# Multiple acyl-CoA dehydrogenase deficiency kills *Mycobacterium tuberculosis* in vitro and during infection

**DOI:** 10.1101/2021.04.19.440201

**Authors:** Tiago Beites, Robert S Jansen, Ruojun Wang, Adrian Jinich, Kyu Rhee, Dirk Schnappinger, Sabine Ehrt

**Affiliations:** Department of Microbiology and Immunology, Weill Cornell Medical College, New York, NY 10065 USA; Division of Infectious Diseases, Department of Medicine, Weill Cornell Medical College, New York, NY, 10065, USA

## Abstract

The human pathogen *Mycobacterium tuberculosis* (Mtb) devotes a significant fraction of its genome to fatty acid metabolism. Although Mtb depends on host fatty acids as a carbon source, fatty acid β-oxidation is mediated by genetically redundant enzymes, which has hampered the development of antitubercular drugs targeting this metabolic pathway. Here, we identify *rv0338c*, referred to as *etfD*_*Mtb*_, to encode a membrane dehydrogenase essential for fatty acid β-oxidation in Mtb. An *etfD* deletion mutant (Δ*etfD*) was incapable of growing on fatty acids in vitro, with long-chain fatty acids being bactericidal, and failed to grow and survive in mice. The Δ*etfD* metabolome revealed a block in β-oxidation at the step catalyzed by acyl-CoA dehydrogenases (ACADs). In many organisms, including humans, ACADs are functionally dependent on an electron transfer flavoprotein (ETF) and cognate dehydrogenase. Immunoprecipitation identified EtfD in complex with FixA (EtfB_Mtb_). FixA (EtfB_Mtb_) and FixB (EtfA_Mtb_) are homologous to the human ETF subunits. Our results demonstrate that EtfBA_Mtb_ constitutes Mtb’s ETF, while EtfD_Mtb_, although not homologous to human EtfD, functions as the dehydrogenase. These findings identify Mtb’s fatty acid β-oxidation as a novel potential target for TB drug development.

## MAIN

Maintenance of an energized membrane is essential for *Mycobacterium tuberculosis* (Mtb) to grow and survive periods of non-replicating persistence^1^. This need has driven tuberculosis (TB) drug development efforts towards Mtb’s energy metabolism. These efforts are supported by the first anti-TB drug approved in over 40 years - the ATP synthase inhibitor bedaquiline^2^.

Mtb’s energy related pathways exhibit varying degrees of vulnerability to inhibition. Uptake of its main carbon sources *in vivo* is performed by specialized transporters; the multi-subunit Mce1 complex transports fatty acids^3^ and the Mce4 complex facilitates the uptake of cholesterol^4^. Inactivation of Mce1 can reduce intracellular growth^5^, while Mce4 was conclusively shown to be essential for survival during the chronic phase of infection^4^. LucA, which acts as a regulator of both Mce1 and Mce4, is also required for wild type levels of Mtb virulence^3^, further supporting that inhibiting the ability to import host lipids affects Mtb’s pathogenicity.

Cholesterol degradation yields multiple products, including acetyl-CoA, propionyl-CoA, succinyl-CoA and pyruvate that can be used for energy generation or lipid biosynthesis^6^. Deletion of genes encoding cholesterol oxidation enzymes attenuated growth^7^ and caused survival defects of Mtb in mice^8,9^. A screen against Mtb residing in the phagosomes of macrophages identified several compounds that target cholesterol metabolism^10^. In contrast, fatty acids are degraded solely through β-oxidation. Mtb’s genome encodes multiple enzymes for each step of β-oxidation, including thirty-four putative acyl-CoA ligases, thirty-five putative acyl-CoA dehydrogenases, twenty-two putative enoyl-CoA dehydratase, five putative β-hydroxyacyl-CoA dehydrogenase and six putative thiolases^11^. Some of these enzymes are necessary for infection, but they also have been shown to play roles in other pathways, such as complex lipid biosynthesis^12^ and cholesterol degradation^13^. Due to the apparent redundancy of Mtb’s fatty acid β-oxidation machinery, this metabolic pathway has thus been presumed to be invulnerable to chemical inhibition.

In the present study, we define an enzyme complex that has previously not been recognized as required for fatty acid degradation in Mtb. It consists of an electron transfer flavoprotein composed by two subunits - FixA (Rv3029c) and FixB (Rv3028c) – and a membrane dehydrogenase (Rv0338c), which we propose to re-name as EtfB_Mtb_, EtfA_Mtb_ and EtfD_Mtb_ based on their human counterparts. Deletion of Mtb’s EtfD causes multiple acyl-CoA dehydrogenase deficiency, which prevents utilization of fatty acids as carbon sources and can kill Mtb in vitro and during mouse infection.

## RESULTS

### A possible role for EtfD_Mtb_ in Mtb’s energy metabolism

EtfD_Mtb_ (Rv0338) is a membrane protein of unknown function predicted to be essential for growth of Mtb on agar plates^14,15^. To experimentally determine its topology, we fused *E. coli* alkaline phosphatase PhoA to EtfD at specific residues of the predicted transmembrane helices. Transport of PhoA outside of the cytoplasm enables reactivity with the chromogenic substrate 5-bromo-4-chloro-3-indolyl phosphate p-toluidine (BCIP). Based on this assay, the soluble portion of EtfD_Mtb_ faces the cytoplasm (Supplementary Fig. 1a), and this topology agrees with the prediction generated by the MEMSAT3 algorithm^16^ (Supplementary Fig. 1b).

Next, we performed in silico analysis, which placed EtfD_Mtb_ into the cluster of orthologue groups 0247 (COG0247) composed of Fe-S oxidoreductases involved in energy production and conversion (Fig. 1a). COG0247 includes a variety of enzymes, including lactate dehydrogenase and the methanogenic-related heterodisulfide reductase; however, most proteins in COG0247 remain uncharacterized and are of unknown function. EtfD_Mtb_ is predicted to be a chimeric enzyme assembled from four domains: an N-terminal domain, which is similar to the gamma subunit of nitrate reductases and putatively binds a cytochrome b; a central 4Fe-4S di-cluster domain similar to succinate dehydrogenases; and two C-terminal cysteine-rich domains (CCG), which are often found in heterodisulfide reductases (Fig. 1b). This analysis led us to hypothesize that EtfD is a component of Mtb’s energy metabolism.

**Fig. 1.**
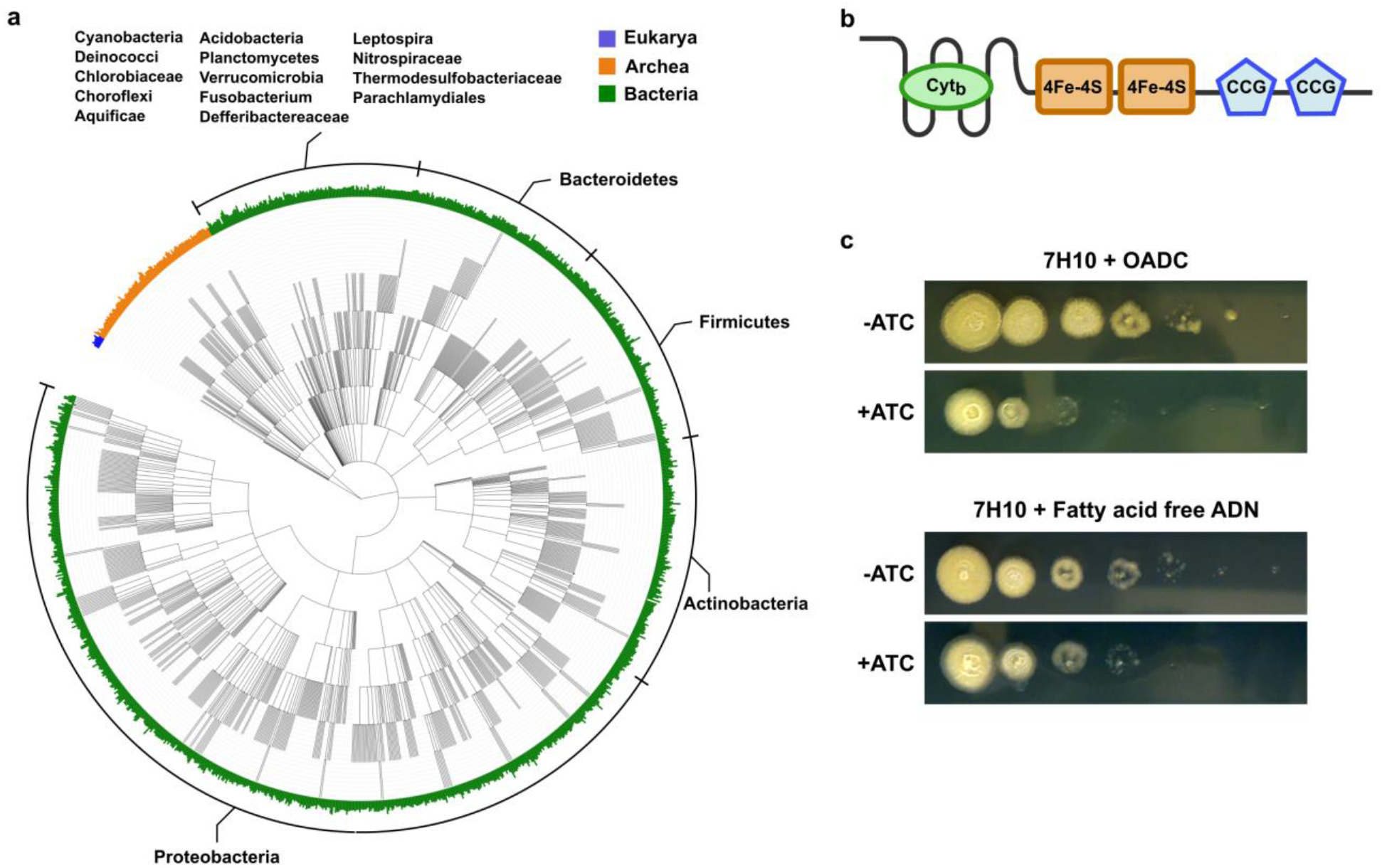
EtfD is a membrane protein possibly involved in Mtb’s energy metabolism. **a**, Rootless phylogenetic tree with species containing members of COG0247 according to the database of the webtool EggNog. The inner colored ring corresponds to life domains, while the outer ring refers to bacteria phyla. Cyt_b_ – cytochrome b, CCG – cysteine-rich domain. **b**, Domain architecture of EtfD based on the algorithms of HHPred and Xtalpred. **c**, Spot assay on solid media with an EtfD-TetOFF strain, where protein levels are controlled by anhydrotetracycline (ATC). Serial dilutions (10^6^ down to 10^1^ bacteria) were incubated for 14 days. These results are representative of 3 independent experiments.

### EtfD_Mtb_ is linked to fatty acid metabolism and it is essential in vivo

To analyze the impact of EtfD_Mtb_ depletion on Mtb’s growth in vitro, we generated a TetOFF strain, in which EtfD levels are controlled by anhydrotetracycline (ATC)-inducible proteolysis^17^. Depletion of EtfD_Mtb_ inhibited Mtb’s growth in regular medium supplemented with oleic acid, albumin, dextrose and catalase (OADC), which is consistent with its predicted essentiality (Fig. 1c). Curiously, when the same medium was supplemented with a fatty acid free enrichment (albumin, dextrose and sodium chloride, ADN), depletion of EtfD_Mtb_ did not impact growth (Fig. 1c). This is similar to the fatty acid sensitive phenotype observed in response to inactivation of Mtb’s type II NADH dehydrogenases^18^ which allowed the genetic deletion of Ndh-2 by growth in a fatty acid free medium. We applied the same strategy to *etfD*_Mtb_ and generated a deletion strain (Δ*etfD*) (Supplementary Fig. 2a). The Δ*etfD*_Mtb_ strain was confirmed by whole genome sequencing (WGS) to not contain additional polymorphisms known to affect growth (Supplementary Table 1). This knockout strain also phenocopied the fatty acid sensitivity observed with the knockdown mutant (Supplementary Fig. 2b).

We next sought to investigate the importance of *etfD*_Mtb_ for pathogenesis in an aerosol model of TB infection in mice. After aerosol infection, Δ*etfD* was unable to grow in mouse lungs and declined in viability from day 14 onwards (Fig. 2a, Supplementary Fig. 3a). In agreement with the CFU data, gross lung pathology showed no lesions in the mice infected with Δ*etfD* (Fig. 2b, Supplementary Fig. 3b). The attenuation was even more pronounced in spleens, where no Δ*etfD* CFU were recovered at any time point (Fig. 2a, Supplementary Fig. 3a). All phenotypes were rescued by reintroducing an intact copy of *etfD*_Mtb_.

**Fig. 2.**
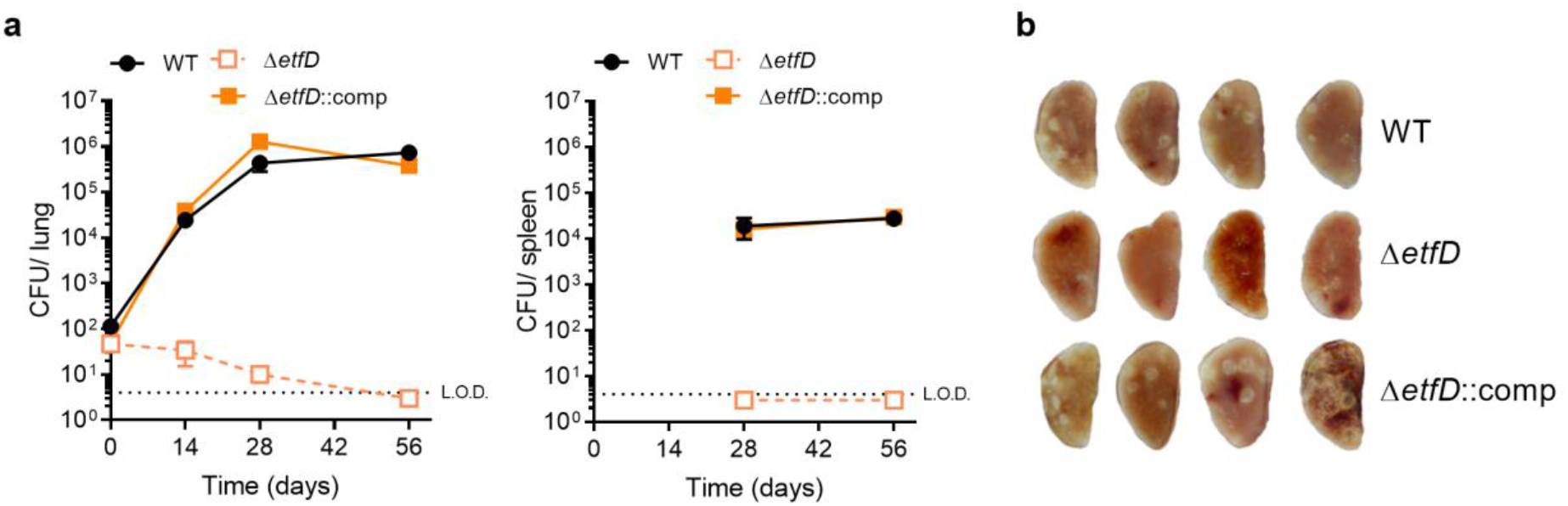
EtfD is essential for growth and survival in vivo. **a**, Growth and persistence of wild type Mtb, Δ*etfD* and the complemented mutant in mouse lungs and spleens. Data are CFU averages from four mice per time point and are representative of two independent experiments. Error bars correspond to standard deviation. “Comp” stands for complemented. L.O.D. stands for limit of detection. **b**, Gross pathology of lungs infected with wild-type Mtb, Δ*etfD* and the complemented mutant at day 56.

### Mtb requires EtfD_Mtb_ to consume fatty acids as a carbon source and to prevent toxicity of long-chain fatty acids

The fatty acid sensitivity of Δ*etfD* suggested that this protein is, directly or indirectly, required for fatty acid metabolism. To test this further, we grew strains in media with different single carbon sources. Δ*etfD* was able to grow in both glycolytic (glycerol) and gluconeogenic (acetic acid and propionic acid) carbon sources, although at a slower rate than the wild type (Fig. 3a). However, longer chain fatty acids (butyric acid, palmitic acid and oleic acid) did not support detectable growth of Δ*etfD* (Fig. 3b). Fatty acids with four carbons or more in length require functional β-oxidation to be utilized, hence these results indicated that Δ*etfD* might display a defect in β-oxidation.

**Fig. 3.**
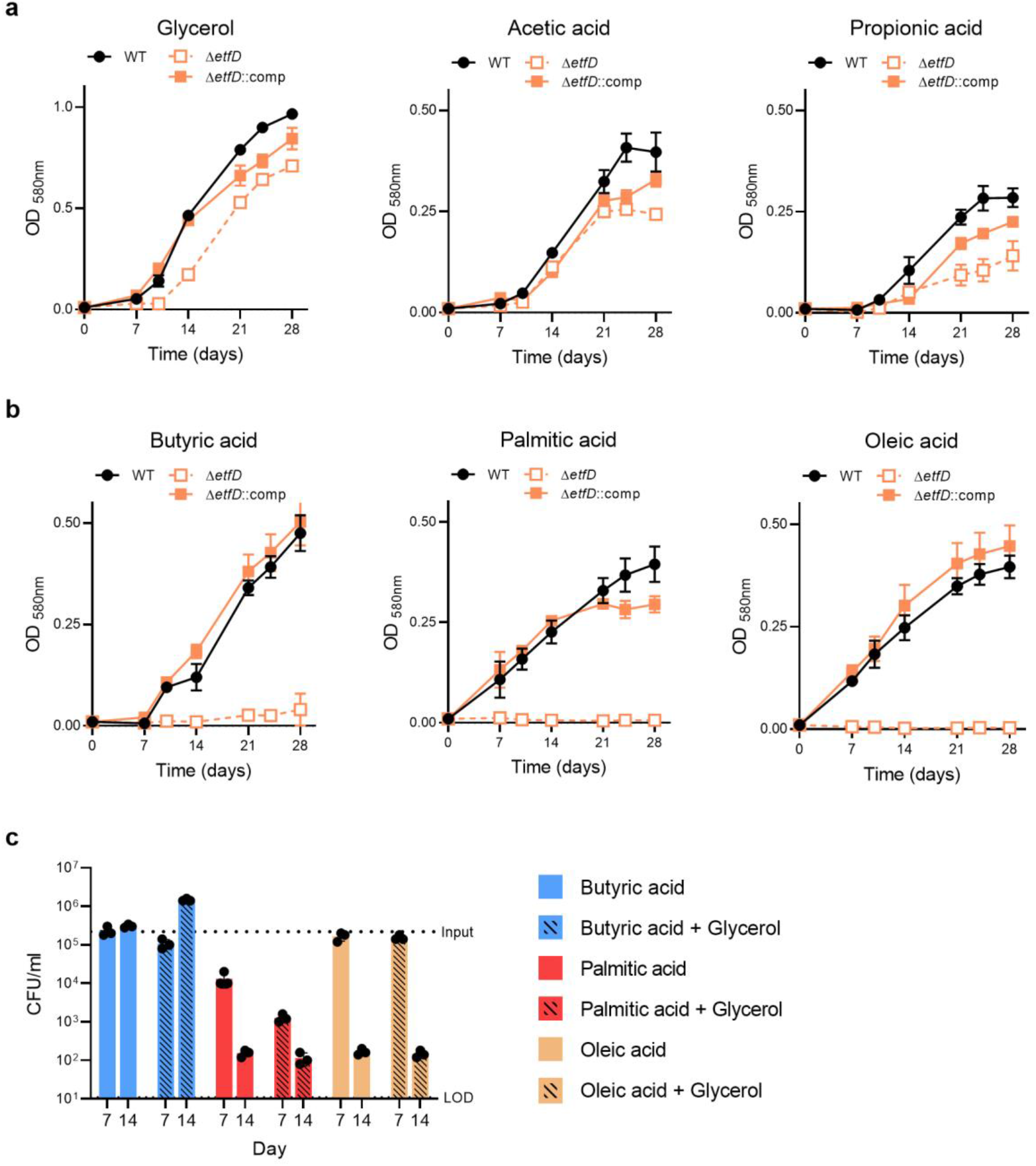
EtfD is necessary for the utilization of fatty acids that require β-oxidation. Strains were grown in media with single carbon sources that (**a**) do not require and (**b**) require β-oxidation for catabolism. Carbon sources were used at the following final concentrations: glycerol 25 mM, acetic acid 25 mM, propionic acid 2.5 mM, butyric acid 2.5 mM, palmitic acid 250 μM and oleic acid 250 μM. To sustain growth and avoid toxicity palmitic acid and oleic acid were replenished every 3 to 4 days for the first 14 days of culture. Data are averages of 3 replicates and are representative of 3 independent experiments. Error bars correspond to standard deviation. “Comp” stands for complemented. **c**, Viability of Δ*etfD* in media with single carbon sources (butyric acid, palmitic acid and oleic acid), or in mixed carbon sources (fatty acids and glycerol) at the same concentrations used in (a) and (b) was assessed at days 7 and 14. Data are averages of 3 replicates and are representative of 3 independent experiments. Error bars correspond to standard deviation. “Comp” stands for complemented. LOD stands for limit of detection.

To distinguish between two possible interpretations, namely 1) Δ*etfD* simply failed to use fatty acids as carbon source and 2) Δ*etfD* was intoxicated by fatty acids, we tested if glycerol could rescue the impaired growth of Δ*etfD* with fatty acids. Glycerol was able to restore growth in medium with butyric acid as carbon source, but it was not able to restore growth with long-chain fatty acids (Supplementary Fig. 4). This argued for a toxic effect and led us to evaluate the impact of fatty acids on the viability of Δ*etfD*. We found that butyric acid is bacteriostatic, while long-chain fatty acids are bactericidal to Δ*etfD* (Fig. 3c). Glycerol did not rescue the bactericidal effect of long-chain fatty acids.

### Mtb acyl-CoA dehydrogenase activity requires EtfD_Mtb_

We applied metabolomics to understand why the consumption of fatty acids is prevented in the absence of EtfD_Mtb_. We focused these studies on ^13^C_4_-labelled butyric acid to isolate the impact of *etfD* disruption on fatty acid consumption from potentially additional confounding effects associated with the toxicity of longer chain fatty acids. Inspection of labelled metabolites in central carbon metabolism confirmed the presence of butyric acid in WT, Δ*etfD*, and the complemented mutant (Fig. 4, Supplementary Fig. 5). All three strains were thus able to import ^13^C_4_-labelled butyric acid.

**Fig. 4.**
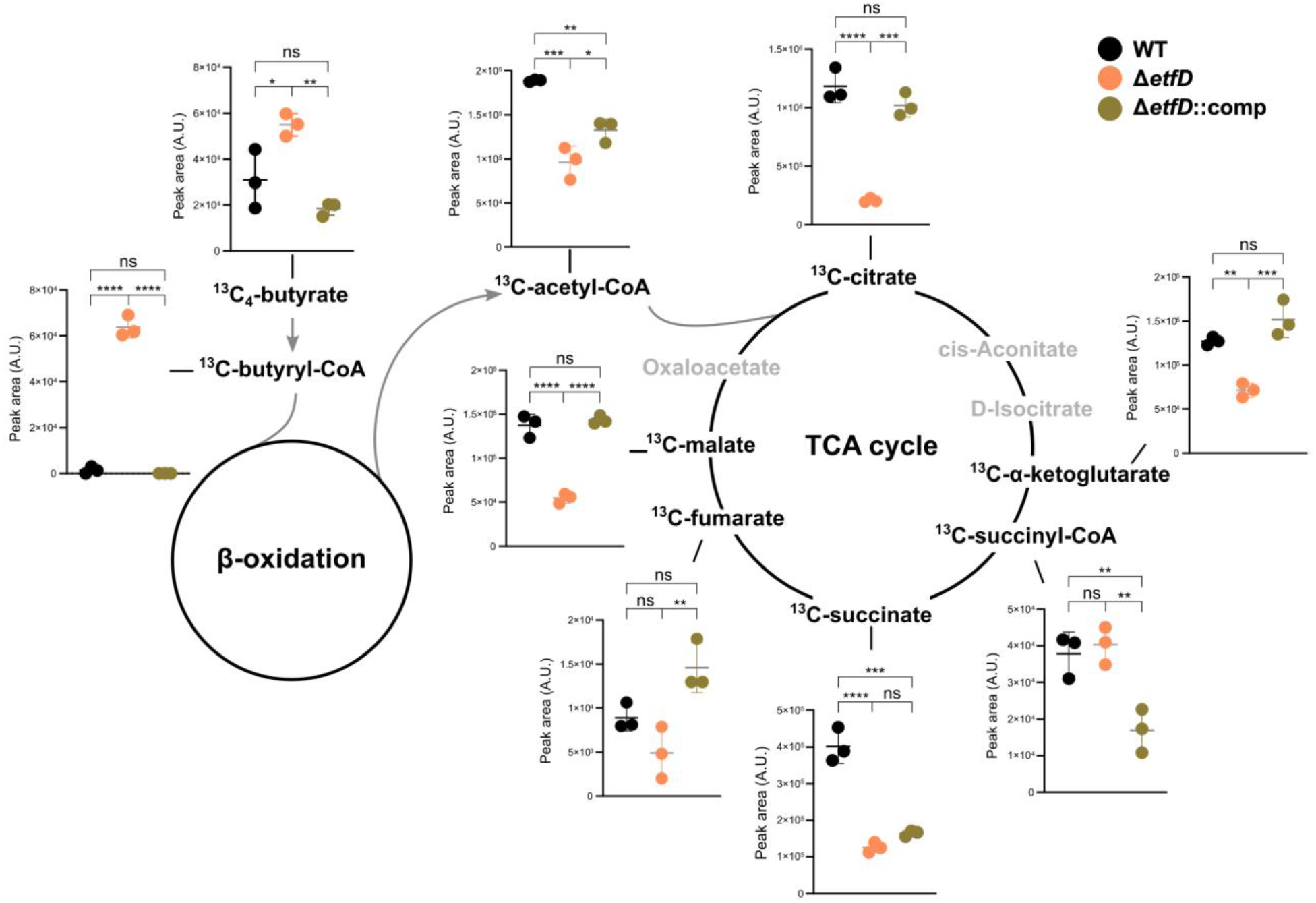
Stable isotope tracing reveals a block in β-oxidation at the level of acyl-CoA dehydrogenases. Strains were grown on filters on top of solid medium permissible to Δ*etfD* growth for 7 days and then transferred to solid media with ^13^C_**4**_-labelled butyric acid (2.5 mM) as single carbon source for 24 hrs. Levels of the indicated ^**13**^C-labeled metabolites (total ^13^C pool, except for ^**13**^C_**4**_-butyrate) were quantified by LC-MS analysis. Data correspond to 3 replicates and are representative of 2 independent experiments. Error bars correspond to standard deviation. “Comp” stands for complemented. Statistical significance was assessed by one-way ANOVA followed by post hoc test (Tukey test; GraphPad Prism). *P < 0.05; **P < 0.01; ***P < 0.001; ****P < 0.0001. ns – not significant. # - metabolites with no statistically significant difference between wild-type and Δ*etfD* in the second independent experiment.

To be catabolized through β-oxidation, butyric acid needs to be transformed by an acyl-CoA ligase into butyryl-CoA. Strikingly, butyryl-CoA accumulated in Δ*etfD* approximately 45-fold relative to WT and the complemented mutant. The remaining intermediates of butyric acid β-oxidation were not detectable in any of the strains, but the pool size of the end-product acetyl-CoA was approximately 2-fold lower in Δ*etfD* than in WT. Labelled TCA cycle intermediates, with the exception of succinyl-CoA, were also partially depleted in Δ*etfD* (Fig. 4, Supplementary Fig. 5). The metabolomic profile of *ΔetfD*, specifically the accumulation of butyryl-CoA, strongly suggested that inactivation of EtfD interfered with the function of acyl-CoA dehydrogenases (ACAD), which in turn impaired fatty acid catabolism.

### EtfD_Mtb_ interacts with an electron transfer protein

We immunoprecipitated EtfD_Mtb_ and identified putative interacting proteins by mass spectrometry. Among the 49 total hits (Supplementary Table 2), 41 were located or predicted to be located at the membrane/cell wall, while the remaining 8 were cytoplasmic proteins. To investigate possible links to β-oxidation – a cytoplasmic process – we focused on the cytoplasmic interactors, which consisted of two flavoproteins EtfB_Mtb_ and Rv1279, the sigma factor SigA, a putative helicase Rv1179c, an exopolyphosphatase Ppx1, the phthiocerol dimycocerosate (PDIM) biosynthesis enzyme PpsC, the protease ClpP2, and Rv1215c, a protein with putative proteolytic activity (Fig. 5a). We were especially interested in the flavoprotein EtfB_Mtb_, because of its homology (30% identity; 87% coverage) with the beta-subunit of the human electron transfer flavoprotein (ETF). Moreover, *etfB*_Mtb_ (annotated as *fixA* – *rv3029c*) forms an operon with *etfA*_Mtb_ (annotated as *fixB* – *rv3028c*), which shares homology with the alpha subunit of the human ETF (41% identity: 98% coverage). In humans, ETF^19^ interacts with a cognate membrane dehydrogenase^20^ (EtfD) and both are required to re-oxidize the FAD co-factor of multiple ACADs. This led us hypothesize that Mtb might display a similar activity. Although EtfD_Mtb_ is not a homologue of the human EtfD, our working model predicted that EtfBA_Mtb_ and EtfD_Mtb_ constitute a complex necessary for the activity of ACADs in Mtb (Fig. 5b).

**Fig. 5.**
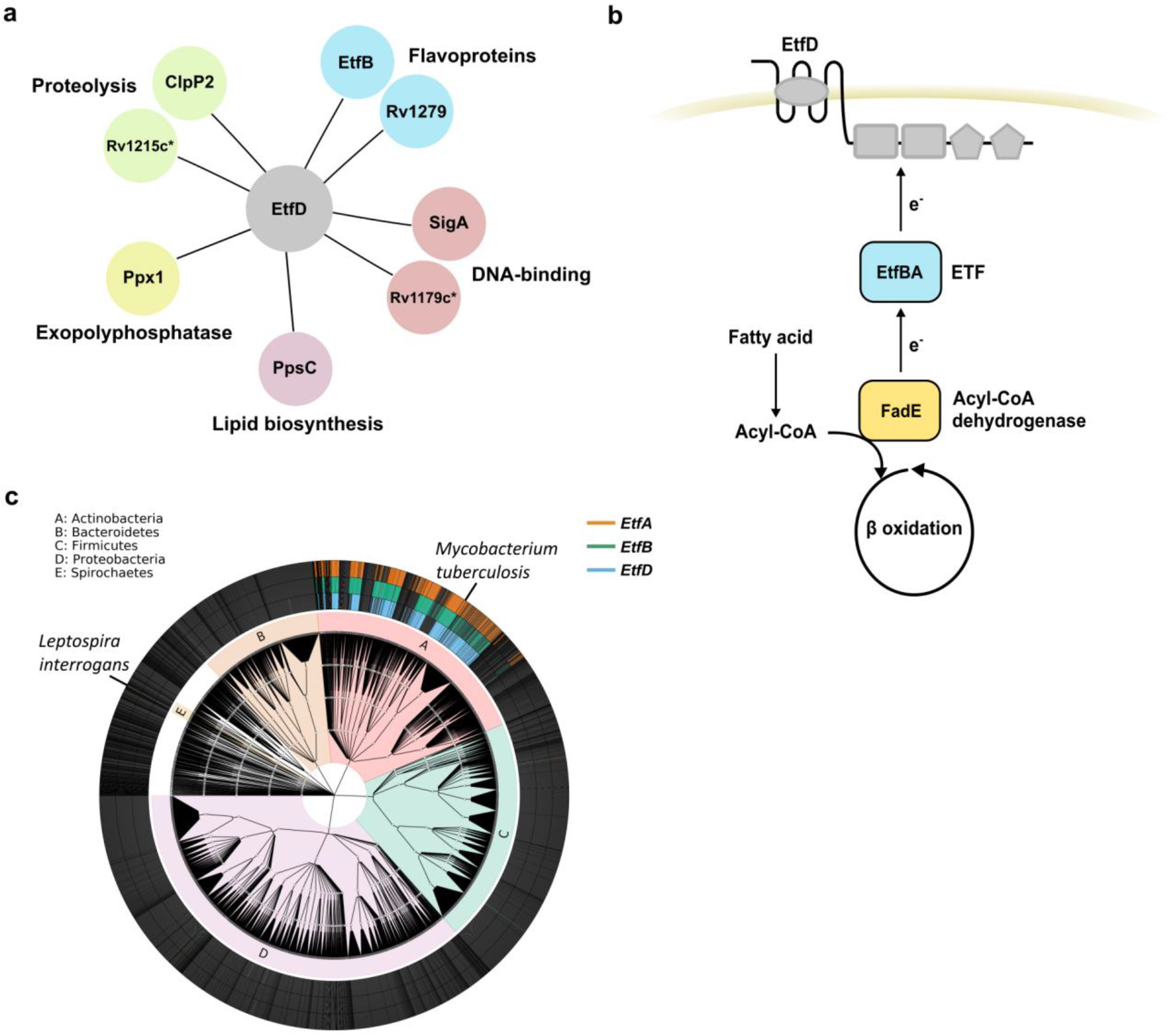
EtfD and EtfBA interaction and co-occurrence. **a**, EtfD cytoplasmic interactors identified by protein co-immunoprecipitation. **b**, Model for the pathway constituted by EtfD and EtfBA. **c**, Uniprot bacterial proteome database was surveyed for EtfD and EtfBA putative homologues. The inner ring corresponds to Phyla and the outer rings represents trains with a hit (>30 % identity and >75% coverage) for EtfD, EtfB or EtfA.* in silico prediction.

To further assess a functional connection between EtfD_Mtb_ and EtfBA_Mtb_, we asked if these proteins co-occur across bacterial proteomes. A BLASTp search against a database of 6240 bacterial proteomes (identity cutoff of >30 % and coverage cutoff of >75%) identified 469 EtfD_Mtb_, 473 EtfB_Mtb_ and 472 EtfA_Mtb_ homologues, 98% of which occur in actinobacteria, with the spirochete *Leptospira interrogans* – the causative agent of leptospirosis – as a notable exception (Figure 5c; Supplementary Data 1). EtfD, EtfB and EtfA showed a strong co-occurrence (p-value <10^−10^), which was suggestive of a functional connection. Curiously, similar to Mtb, *L. interrogans* is proposed to use host-derived fatty acids as the primary carbon sources during infection^21^.

These results support the hypothesis that EtfD_Mtb_ serves as a dehydrogenase for EtfBA_Mtb_, which in turn is necessary for the activity of ACADs.

### EtfBA_Mtb_ and EtfD_Mtb_ are required for acyl-CoA dehydrogenase activity

If EtfBA_Mtb_ and EtfD_Mtb_ participate in the same biochemical pathway then inactivation of EtfBA_Mtb_ should also impair the of use fatty acids as single carbon sources by Mtb. To test this prediction, we first isolated a knockout strain for *etfBA* in fatty acid free medium (Supplementary Fig. 6) and confirmed its genetic identity through WGS (Supplementary Table 1). We then grew wild type, Δ*etfBA* and complemented Δ*etfBA* in media with different carbon sources. Δ*etfBA* was able to grow with glycerol, albeit slower than WT (Fig. 6a). In contrast, we did not detect growth in either butyric acid or oleic acid as single carbon sources, corroborating our prediction (Fig. 6b and c).

**Fig. 6.**
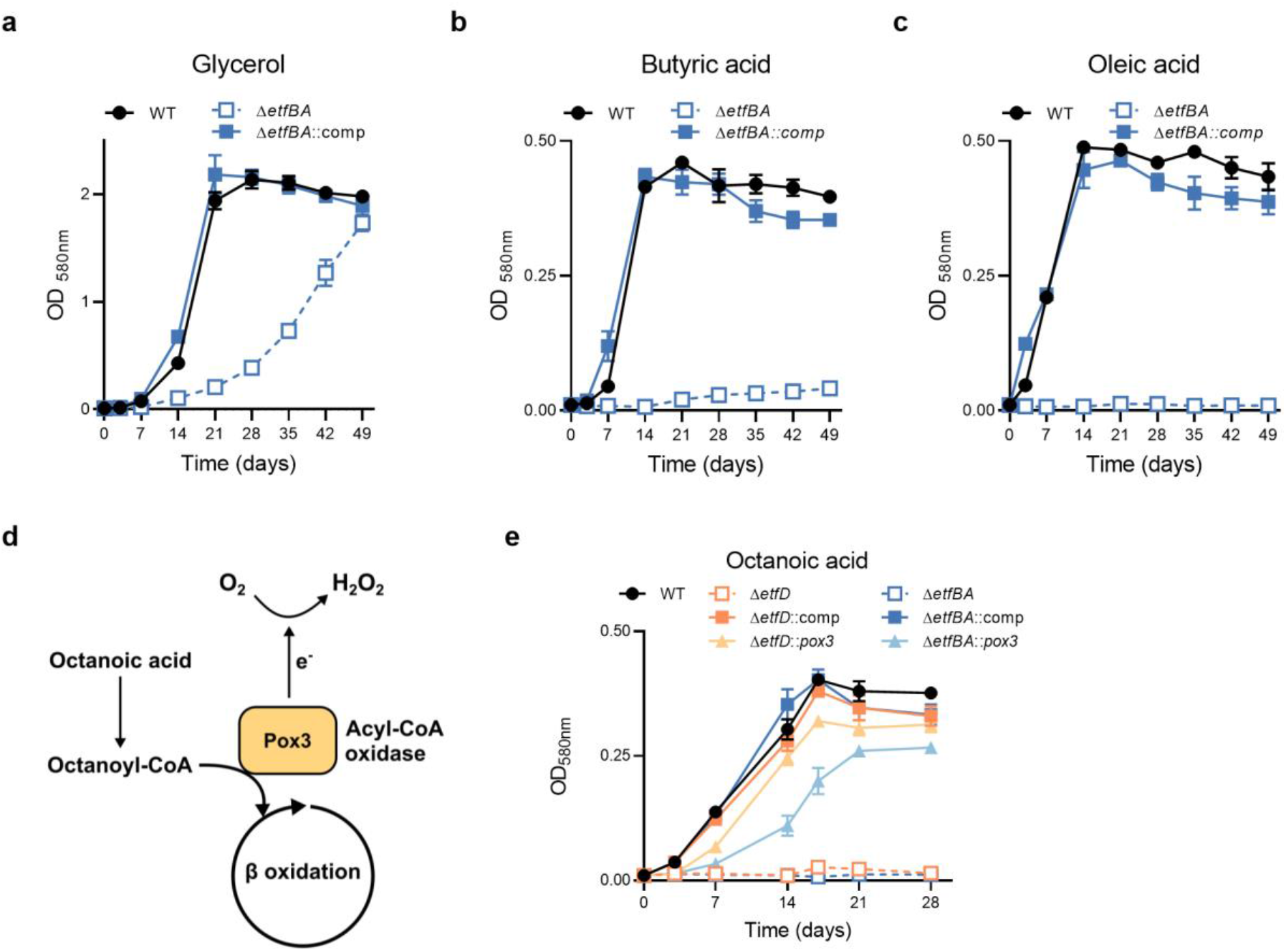
EtfD and EtfBA constitute a pathway necessary for the activity of acyl-coA dehydrogenases. Growth with 25 mM glycerol (**a**), 2.5 mM butyric acid (**b**) and 250 μM oleic acid (**c**) as single carbon sources. Oleic acid was replenished every 3 to 4 days for the first 14 days of culture to support growth and minimize toxicity. Data are averages of 3 replicates and are representative of 3 independent experiments. Error bars correspond to standard deviation. “Comp” stands for complemented. **d**, Diagram representing acyl-coA oxidase activity on octanoic acid. **e**, Rescue of Δ*etfD* and Δ*etfBA* growth with 250 μM octanoic acid as sole carbon source with the expression of the acyl-CoA oxidase Pox3. Octanoic acid was replenished every 3 to 4 days for the first 14 days of culture to support growth and minimize toxicity. Data are averages of 3 replicates.

To independently test the role of EtfBA_Mtb_ and EtfD_Mtb_ in fatty acid oxidation, we assessed if an acyl-CoA oxidase (ACO) could allow both Δ*etfBA* and Δ*etfD* to grow with fatty acids. ACOs are peroxisomal enzymes that catalyze the same reaction as ACADs using molecular oxygen to re-oxidize FAD rather than ETF/ETFD^22^. The gene *pox3*, which encodes the extensively characterized ACO Pox3 from the yeast *Yarrowia lipolytica*^23,24^ (Fig. 6d), was codon adapted for Mtb and expressed under the control of a strong, constitutive promoter in both Δ*etfBA* and Δ*etfD*. Pox3 has a high affinity for fatty acids with six and eight carbons chain length and low affinity for most other fatty acids^23^; thus, we grew strains in a medium with octanoic acid (high affinity), butyric acid (low affinity) or oleic acid (low affinity) as single carbon sources. Δ*etfD* was not able to utilize octanoic acid as single carbon source, further confirming its inability to oxidize multiple fatty acids, while the growth rate of Δ*etfD*::*pox3* was similar to that of WT and complemented strain (Fig. 6e). A similar result was obtained with butyric acid, although it took longer for Δ*etfD*::*pox3* to reach wild type density (Supplementary Fig. 7). As expected, based on the affinity profile and function of Pox3, *pox3* did not rescue Δ*etfD* growth with oleic acid as single carbon source and it did not alter the growth rate on glycerol. Similarly, Δ*etfBA* did not grow on octanoic acid as single carbon source, while Δ*etfBA*::*pox3* was able to grow in the same medium, although reaching a lower final optical density than WT and complemented mutant. Δ*etfBA*::*pox3* was able to grow with butyric acid entering stationary phase 21 days after wild-type and complemented strain, it did not grow with oleic acid and it conferred a slight advantage on glycerol when compared with Δ*etfBA* (Fig. 6e and Supplementary Fig. 7).

These results confirmed that EtfBA_Mtb_ and EtfD_Mtb_ constitute a complex necessary for the activity of fatty acid β-oxidation ACADs.

## DISCUSSION

Mtb’s energy metabolism displays a remarkable plasticity which supports its adaptation to a multitude of host microenvironments^25^. Its unusual domain structure suggested that the membrane protein encoded by *rv0338c* could be an unrecognized, essential component of Mtb’s energy metabolism. Our studies identified Rv0338c as a member of a short electron transfer pathway essential for ACADs activity. Based on the similarity of this system to the human ETF system we propose to rename the respective Mtb genes as *etfD*_Mtb_ (*rv0338c*), *etfB*_Mtb_ (*rv3028c*) and *etfA*_Mtb_ (*rv3029c*).

The increased susceptibility to fatty acids of an EtfD_Mtb_ TetOff strain suggested a possible connection with fatty acid metabolism. Accordingly, our data showed that an *etfD*_Mtb_ deletion mutant was not capable of utilizing fatty acids with four carbons or more as single carbon sources, thus indicating impairment in β-oxidation. This was associated with different outcomes regarding viability: butyric acid (short-chain) was bacteriostatic, while palmitic acid and oleic acid (long-chain) were bactericidal. Importantly, Δ*etfD* did not grow and presented a survival defect in mice. Long-chain fatty acids, including oleic acid, are common components of human macrophages^26^ and a carbon source for Mtb. Hence, the inability to utilize fatty acids together with the increased susceptibility to long-chain fatty acids likely explain the in vivo essentiality of EtfD_Mtb_. The mechanism of long-chain fatty acid toxicity to Mtb is still poorly understood, even though it was first described in the 1940’s^27^. In other bacteria, several mechanistic explanations have been proposed for the bactericidal activity of long-chain fatty acids, including membrane potential disruption^28,29^, oxidative stress induction^30^, or fatty acid biosynthesis inhibition^31^.

The metabolome of Δ*etfD* in medium with butyric acid as sole carbon source revealed an accumulation of butyryl-CoA which indicated an impairment in β-oxidation. This can explain the inability of Δ*etfD* to oxidize fatty acids and connects EtfD_Mtb_ with the β-oxidation enzymes that act on acyl-CoAs – the ACADs. None of Mtb’s thirty-five annotated ACADs are essential in vivo^32^ and there are no reports showing a specific ACAD being essential in vitro for the utilization of any fatty acid. Thus, ACADs are likely to be functionally redundant. However, this redundancy was not sufficient to support growth of Δ*etfD* in media with fatty acids as single carbon sources, strongly suggesting that EtfD_Mtb_ is necessary for the activity of multiple, if not all, ACADs.

Immunoprecipitation using EtfD_Mtb_ as bait revealed multiple possible interacting proteins, suggesting that EtfD_Mtb_ might integrate several pathways in Mtb. We were especially interested in the interaction with the cytoplasmic protein EtfB_Mtb_, which together with EtfA_Mtb_ constitute a putative electron transfer flavoprotein (ETF). The human homologues (EtfAB) form an enzyme that re-oxidizes the FAD co-factor of multiple ACADs and transfers the electrons to the membrane bound oxidoreductase electron transfer flavoprotein dehydrogenase (EtfD), which then reduces the electron carrier ubiquinone, hence contributing to the generation of energy^33^. Mutations rendering defects in EtfAB or EtfD lead to a metabolic disease named multiple acyl-CoA dehydrogenase deficiency, which among other outcomes is characterized by the inability to oxidize fatty acids^34^. This led us to hypothesize that Mtb’s EtfBA_Mtb_ and EtfD_Mtb_ might work together in a similar pathway. That EtfD_Mtb_ and EtfBA_Mtb_ show a strong pattern of co-occurrence across bacterial proteomes and Δ*etfBA* was unable to utilize fatty acids as carbon sources were strong indications in favor of our hypothesis. Nevertheless, for a yet unclear mechanism, it was notable that Δ*etfBA* grew more slowly than Δ*etfD*. The expression of the acyl-CoA oxidase Pox3^23,24^, an enzyme that catalyzes the same reaction as ACADs, but uses molecular oxygen as an electron acceptor^22^, was able to rescue the growth of both Δ*etfD* and Δ*etfBA* in fatty acids as single carbon source, showing that in both cases β-oxidation was impaired at the ACAD step and confirming our proposed hypothesis. Although we report a function in β-oxidation, it is possible that these proteins may have additional roles in Mtb. It has recently been reported that both EtfBA_Mtb_ and EtfD_Mtb_ are essential for resisting toxicity in media containing heme^35^. This further strengthened the functional relation between these proteins and suggested a role in iron metabolism. Interestingly, transcript levels of *etfD*_Mtb_, but not *etfBA*_Mtb_, respond to the iron content of the medium in an IdeR (iron metabolism transcriptional regulator) dependent manner^36,37^. Whether the EtfBA_Mtb_/ EtfD_Mtb_ complex has a direct or indirect role in heme utilization and if this has any in vivo relevance remain to be addressed.

In conclusion, we have identified a complex composed of EtfBA_Mtb_ and the cognate membrane dehydrogenase EtfD_Mtb_ that is required for the function of multiple ACADs. This complex constitutes a previously unsuspected vulnerable component of Mtb’s β-oxidation machinery. EtfD_Mtb_ has no structural homologues in humans and was recently proposed as the target of a compound that is bactericidal against Mtb^38^, which suggests that it could serve as a novel target for TB drug development. The presence of EtfD_Mtb_ and EtfBA_Mtb_ homologues in *Leptospira interrogans* suggests that this pathway might be relevant for other pathogens that rely on host fatty acids as carbon sources^21^.

## METHODS

### Culture conditions

For cloning purposes we used *Escherichia coli* as a host, which was cultured in LB medium at 37 °C. *Mtb* was cultured at 37 °C in different media: Middlebrook 7H9 supplemented with 0.2% glycerol, 0.05% tyloxapol, and ADNaCl (0.5% fatty acid free BSA from Roche, 0.2% dextrose and 0.85% NaCl) or in Middlebrook 7H10 supplemented with 0.5% glycerol and 10% oleic acid-albumin-dextrose-catalase (OADC), and in a modified Sauton’s minimal medium (0.05% potassium dihydrogen phosphate, 0.05% magnesium sulfate heptahydrate, 0.2% citric acid, 0.005% ferric ammonium citrate, and 0.0001% zinc sulfate) supplemented with 0.05% tyloxapol, 0.4% glucose, 0.2% glycerol, and ADNaCl with fatty-acid-free BSA (Roche). Modified Sauton’s solid medium contained 1.5% bactoagar (BD) and glycerol at a higher concentration (0.5%). For single or mixed carbon source cultures we have used glycerol 25 mM, sodium acetate 2.5 mM, propionic acid 2.5 mM and butyric acid 2.5 mM. Octanoic acid, palmitic acid and oleic acid were added at a final concentration of 200 μM and were replenished every 3-4 days for the first 14 days, since these fatty acids are toxic if added initially in the mM range. *Mycobacterium smegmatis* MC^2^ 155 was cultured in Middlebrook 7H10 supplemented with 0.2% glycerol at 37 °C. Antibiotics were used at the following final concentrations: carbenicillin 100 μg/ml, hygromycin 50 μg/ml and kanamycin 50 μg/ml.

### Mutant construction

Mtb H37Rv *etfD*_Mtb_ conditional knockdown was generated using a previously described strategy that control expression through proteolysis^39^. Briefly, a Flag tag and DAS+4 tag were added to the 3’ end of *etfD*_Mtb_, at the 3′ end of the target gene. This strain was then transformed with a plasmid expressing the adaptor protein SspB under the control of TetR regulated promoters. In the presence of anhydrotetracycline (ATC) 500 μg/ml, TetR loses affinity to the promoter and *sspB* expression is derepressed. SspB acts by delivering DAS+4-tagged proteins to the native ClpXP protease. Hence, when ATC is added to the culture, SspB expression is induced and EtfD_Mtb_-DAS-tag is degraded, working as a TetOFF system (EtfD_Mtb_-TetOFF).

We have obtained deletion mutants through recombineering by using Mtb H37Rv expressing the recombinase RecET. Constructs with the hygromycin resistant gene (hygR) flanked by 500 bp upstream and downstream of the target loci were synthesized (GeneScript). In the case of *etfD*, since the flanking gene *aspC* is in the same orientation and it is essential for growth^40^, we have included in the construct the constitutive promoter hsp60 to avoid polar effects. Mutants were selected in modified Sauton’s with hygromycin. The plasmid expressing *recET* was counterselected by growing the deleted mutants in modified Sauton’s supplemented with sucrose 10 %. For complementation we have cloned *etfD* and *etfBA* under the control of the promoter phsp60 into a plasmid with a kanamycin resistant cassette that integrates at the att-L5 site (pMCK-phsp60-*etfD* and pMCK-phsp60-*etfBA*) and transformed the deleted mutants. The gene *pox3* from *Yarrowia lipolytica* was codon adapted for *Mtb* use, synthesized (GeneScript), cloned into a plasmid under the control of the promoter pTB38, with a kanamycin resistant cassette that integrates in the att-L5 site (pMCK-pTB38-*pox3*) and transformed into both *etfD*_Mtb_ and *etfBA*_Mtb_ deletion mutants. All generated strains and plasmids are listed in Supplementary Tables 3 and 4, respectively.

### Whole genome sequencing

The genetic identity of Δ*etfD* and Δ*etfBA* was confirmed by whole genome sequencing (WGS). Between 150 and 200 ng of genomic DNA was sheared acoustically and HiSeq sequencing libraries were prepared using the KAPA Hyper Prep Kit (Roche). PCR amplification of the libraries was carried out for 10 cycles. 5–10 × 106 50-bp paired-end reads were obtained for each sample on an Illumina HiSeq 2500 using the TruSeq SBS Kit v3 (Illumina). Post-run demultiplexing and adapter removal were performed and fastq files were inspected using fastqc (Andrews S. (2010). FastQC: a quality control tool for high throughput sequence data. Available at: http://www.bioinformatics.babraham.ac.uk/projects/fastqc). Trimmed fastq files were then aligned to the reference genome (M. tuberculosis H37RvCO; NZ_CM001515.1) using bwa mem47. Bam files were sorted and merged using samtools48. Read groups were added and bam files de-duplicated using Picard tools and GATK best-practices were followed for SNP and indel detection49. Gene knockouts and cassette insertions were verified for all strains by direct comparison of reads spanning insertion points to plasmid maps and the genome sequence. Reads coverage data was obtained from the software Integrative Genomics Viewer (IGV)^41-43^. Sequencing data was deposited in NCBI’s Sequence Read Archive (SRA) database under the BioProject PRJNA670664.

### PhoA fusion assay

Truncated versions of EtfD_Mtb_ were fused with the *E. coli* alkalyne phosphatase PhoA. This enzyme requires the oxidative environment of the periplasm to be active and degrades the substrate BCIP generating a blue precipitate. We have fused *phoA* at different residues located in the transmembrane domains. Positive control consisted in PhoA fused with the antigen 85B, while the negative control was PhoA alone^44^. All plasmids were transformed in *M. smegmatis* MC2 and the assay was performed in LB plates with and without BCIP.

### Mouse infection

Mouse experiments were performed in accordance with the Guide for the Care and Use of Laboratory Animals of the National Institutes of Health, with approval from the Institutional Animal Care and Use Committee of Weill Cornell Medicine. Female C57BL/6 mice (Jackson Labs) were infected with ∼100–200 CFU/mouse using an Inhalation Exposure System (Glas-Col). Strains were grown to mid-exponential phase and single-cell suspensions were prepared in PBS with 0.05% Tween 80, and then resuspended in PBS. Lungs and spleen were homogenized in PBS and plated on modified Sauton’s medium to determine CFU/organ at the indicated time points.

### Metabolomics

Strains were grown in modified Sauton’s until an OD_580_nm of 1 and 1 ml of culture was used to seed filters^45^ placed on top of solid modified Sauton’s medium. Bacteria grew for 7 days, after which the filters were transferred to solid modified Sauton’s medium with butyric acid or ^13^C-labelled butyric acid (Cambridge Isotope Laboratories, Inc) at a final concentration of 2.5 mM for 24 h. For metabolite extraction bacteria were disrupted by bead beating 3 cycles, 50 seconds (Precellys 24, Bertia technologies) in a solution of acetonitrile:methanol:water (4:4:2).

The relative abundances of butyric acid, CoA species and TCA intermediates were determined using an ion-pairing LC-MS system, as previously described^46^. In brief, samples (5 uL) were injected onto a ZORBAX RRHD Extend-C18 column (2.1 x 150 mm, 1.8 µm; Agilent Technologies) with a ZORBAX SB-C8 (2.1 mm × 30 mm, 3.5 μm; Agilent Technologies) precolumn heated to 40 °C and separated using a gradient of methanol in 5 mM tributylamine/5.5 mM acetate. Post-column, 10% dimethyl sulfoxide in acetone (0.2 mL/min) was mixed with the mobile phases to increase sensitivity. Detection was performed from m/z 50-1100, using an Agilent Accurate Mass 6230 Time of Flight (TOF) spectrometer with Agilent Jet Stream electrospray ionization source operating in the negative ionization mode. Incorporation of ^13^C was quantitated and corrected for natural ^13^C abundance using Profinder B.08.00 (Agilent Technologies). LC-MS data was deposited in the MetaboLights database^47^ under the accession code MTBLS2374 (www.ebi.ac.uk/metabolights/MTBLS2374).

### Immunoprecipitation

We transformed Δ*etfD* with a plasmid expressing flag tagged EtfD under hsp60 promoter (pMEK-Phsp60-etfD_Mtb_-flag) and used WT Mtb expressing only the flag tag as a control. Mtb whole-cell lysates were collected from 120 ml log phase culture in butyric acid single carbon source Sauton’s medium, incubated with 1% DDM for two hours on ice, followed by anti-Flag beads (Sigma) overnight incubation with gentle rotation. Beads were collected on the second day, washed with lysis buffer (50 mM Tris-HCl, 50 mM NaCl, pH 7.4), and eluted with 100 ng/μl Flag peptide. The eluates were resolved on SDS-PAGE before mass spectrometry.

For mass spectrometry analysis, the total spectrum count (TSC) from biological duplicates were summed. We calculated the ratio of summed TSC from EtfD_Mtb_-Flag vs. Flag control and used a cut-off of >=10.

### In silico analysis

Transmembrane domain topology of EtfD_Mtb_ was performed in MEMSAT3^16^. Domain architecture of EtfD_Mtb_, EtfB and EtfA was based on HHPred^48^ and XtalPred^49^. Eggnog ^50^ was used for the cluster of orthologous groups analysis (COG). The members of COG247 that include *etfD* were used to generate a rootless phylogenetic tree in iTOL^51^.

To analyze the presence or absence of EtfD_Mtb_, EtfB_Mtb_, and EtfA_Mtb_ homologues across bacterial species, we obtained the set of UniProt reference bacterial proteomes, which are are selected both manually and algorithmically by UniProt as landmarks in (bacterial) proteome space^52^. We discarded proteomes with no taxonomic labels and performed the analysis on a final set of 6240 bacterial reference proteomes. Using EtfD_Mtb_, EtfB_Mtb_, and EtfA_Mtb_ as query protein sequences, we used the following protein BLAST (BLASTp) parameter values: identity cutoff of >30%, coverage cutoff of >75%, e-value cutoff of 10-3. Visual representations of phylogenies with surrounding color-coded rings were generated using the software tool GraPhLan^53^, with the phylogenetic try built from the taxonomic categorization of the 6240 UniProt bacterial reference proteomes. To evaluate the statistical significance of co-occurrence of EtfD_Mtb_ and EtfBA_Mtb_ in Actinobacteria, we performed a hypergeometric test to evaluate the probability of observing k species with EtfD and EtfBA homologues, given M Actinobacterial species, n species with EtfD homologues, and N species with EtfBA homologues.

### Quantification and statistical analysis

Generation of graphics and data analyses were performed in Prism version 8.0 software (GraphPad).

## ACKNOWLEDGEMENTS

We thank J. McConnell and C. Trujillo for help with strain generation and in vivo characterization. We thank K.G. Papavinasasundaram (University of Massachusetts), R. Aslebagh and S.A. Shaffer (University of Massachusetts, Mass Spectrometry Facility) for LC-MS/MS analysis. We acknowledge J.M. Bean from MSKCC and the use of the Integrated Genomics Operation Core at MSKCC, funded by the NCI Cancer Center Support Grant (CCSG, P30 CA08748), Cycle for Survival and the Marie-Josée and Henry R. Kravis Center for Molecular Oncology. D.S and S.E were supported by Tri-Institutional TB Research Unit U19AI111143. T.B. was supported by a Potts Memorial Foundation fellowship.

## Author contributions

S.E. and D.S. conceived ideas, supervised the study and revised the manuscript. T.B. conceived ideas, performed experimental work, analysed and interpreted data and wrote the manuscript. R.S.J., R.W. performed experimental work, analysed and interpreted data and reviewed the manuscript. A.J. performed the in silico analysis and reviewed the manuscript. K.Y.R supervised the metabolomics experiments and reviewed the manuscript.

