## Supplementary Information for "Multiple acyl-CoA dehydrogenase deficiency kills *Mycobacterium tuberculosis* in vitro and during infection"

### Suplementary Information

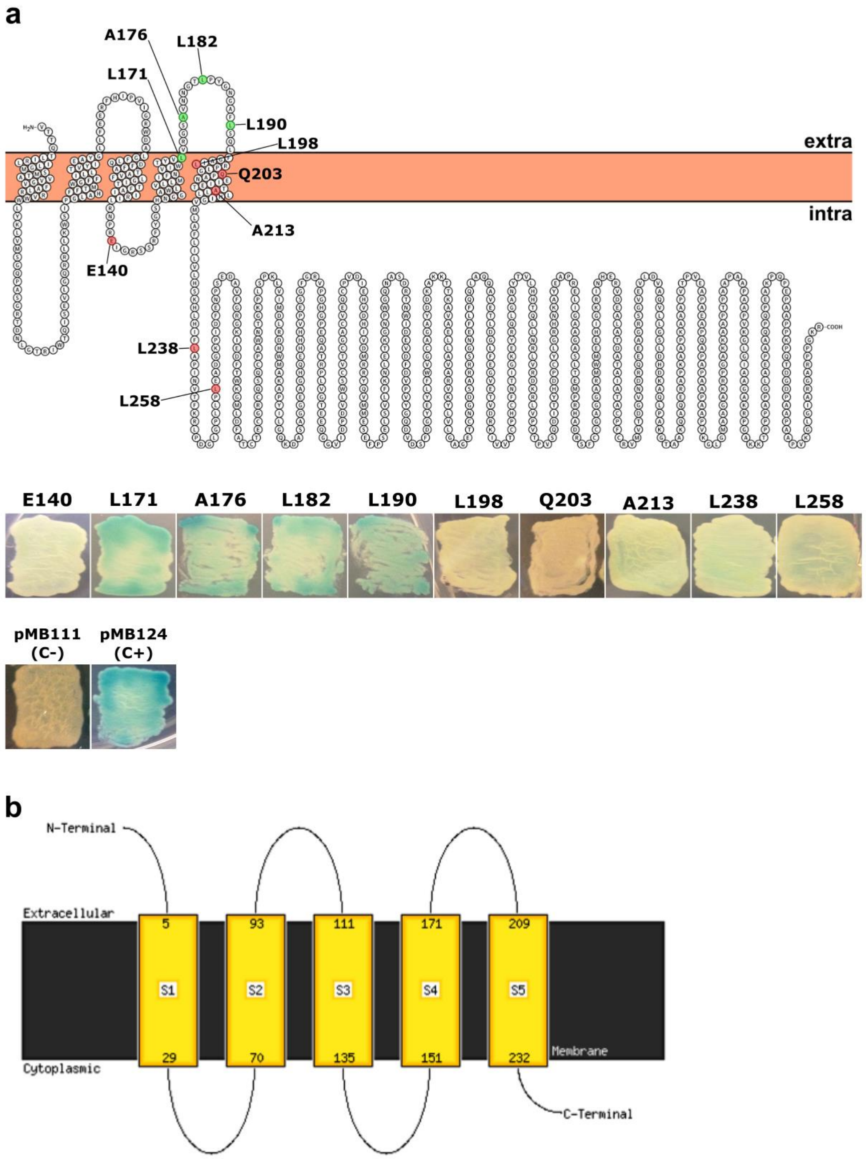


**Supplementary Fig. 1 EtfD has 5 transmembrane domains and the soluble portion faces the periplasm. a**, *E. coli* alkaline phosphatase PhoA was fused at different residues of the EtfD (Rv0338c) transmembrane domain and soluble portion. The constructs were expressed in *Mycobacterium smegmatis* and the strains grew in the presence of the chromogenic PhoA substrate BCIP. As positive control (c+) PhoA is fused with the antigen 85B that contains a secretion signal (PhoA requires the oxidative environment of the periplasm to be active), while the negative control (c-) lacks any secretion signal. These results are representative of 2 independent experiments. **b**, EtfD (Rv0338c) topology prediction with the MEMSAT algorithm.

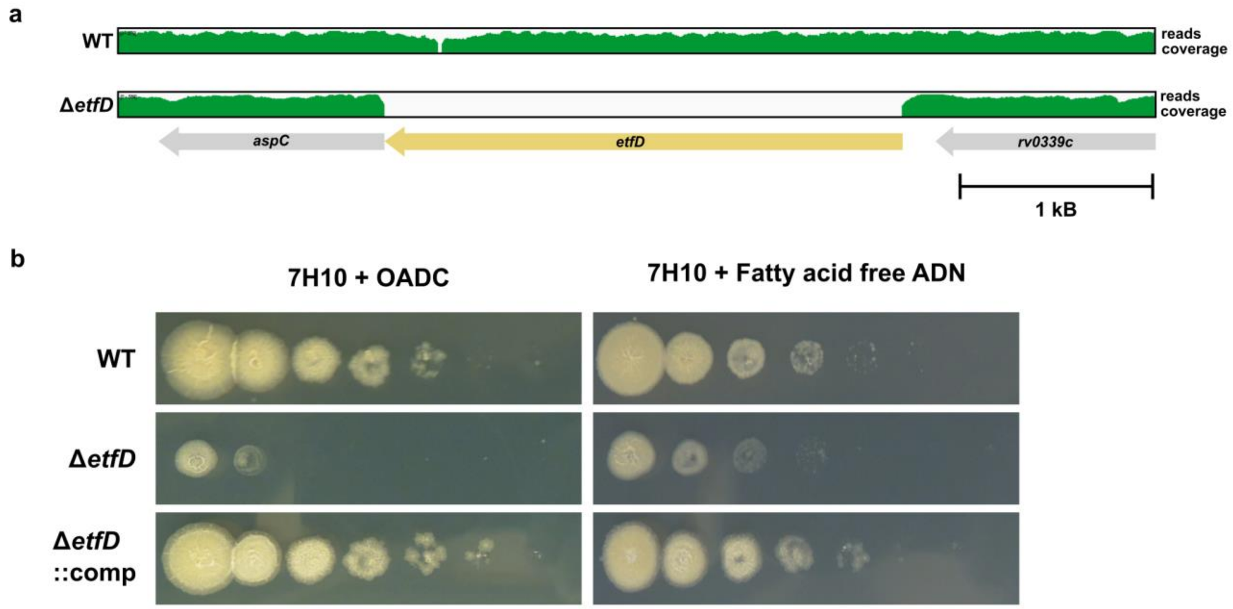


**Supplementary Fig. 2. Δ*etfD* genetic identity confirmation**. **a**, Whole genome sequencing coverage showing the *etfD* wild-type locus and gene deletion in Δ*etfD*. **b,** Spot assay on solid media. Serial dilutions ($10^6$ down to $10^1$ bacteria) were incubated for 14 days. These results are representative of 3 independent experiment.

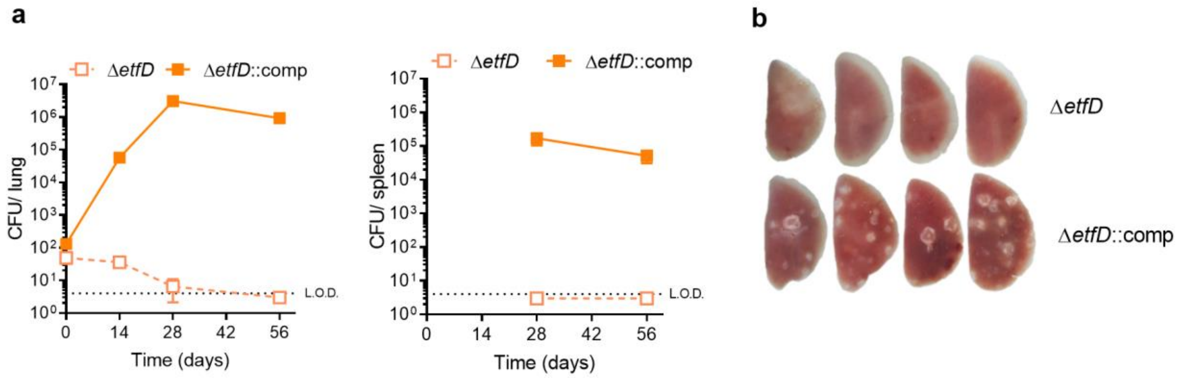


**Supplementary Fig. 3. EtfD is essential for growth and survival in vivo.** Second experiment of growth and persistence of Δ*etfD* in vivo. **a**, Growth and persistence of wild type Mtb, Δ*etfD* and the complemented mutant in mouse lungs and spleens. Data are CFU averages from four mice per time point and are representative of two independent experiments. Error bars correspond to standard deviation. "Comp" stands for complemented. L.O.D. stands for limit of detection. **b**, Gross pathology of lungs infected with wild-type Mtb, Δ*etfD* and the complemented mutant at day 56.

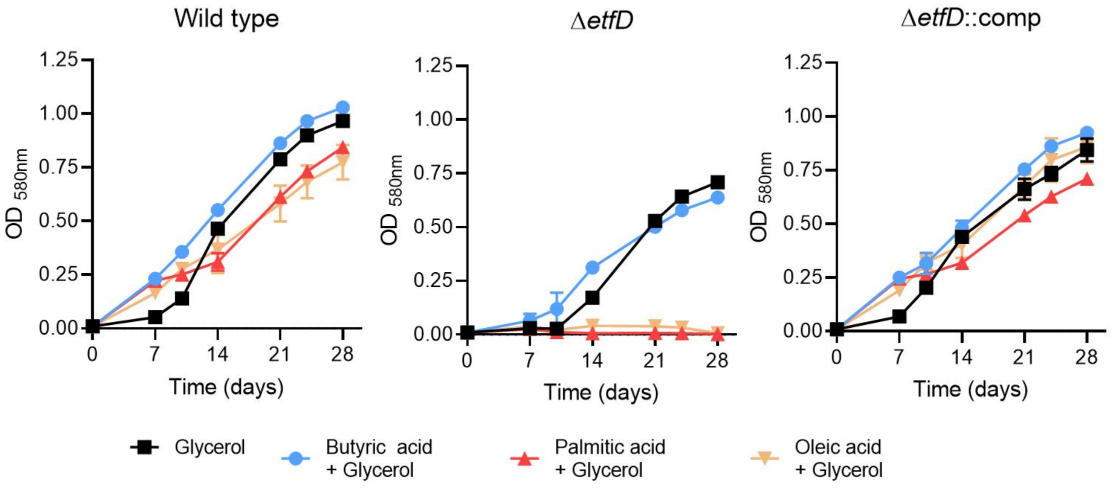


**Supplementary Fig. 4. Long-chain fatty acids are toxic to Δ*etfD.*** Strains were grown in media with mixed carbon sources containing 25 mM glycerol and one of the following fatty acids: 2.5 mM butyric acid, 250 μM palmitic acid and 250 μM oleic acid. Glycerol (25 mM) alone was used as a positive control for Δ*etfD*. Palmitic acid and oleic acid were replenished every 3 to 4 days to sustain growth and avoid toxicity. Data are averages of 3 replicates and are representative of 3 independent experiments. Error bars correspond to standard deviation. "Comp" stands for complemented.

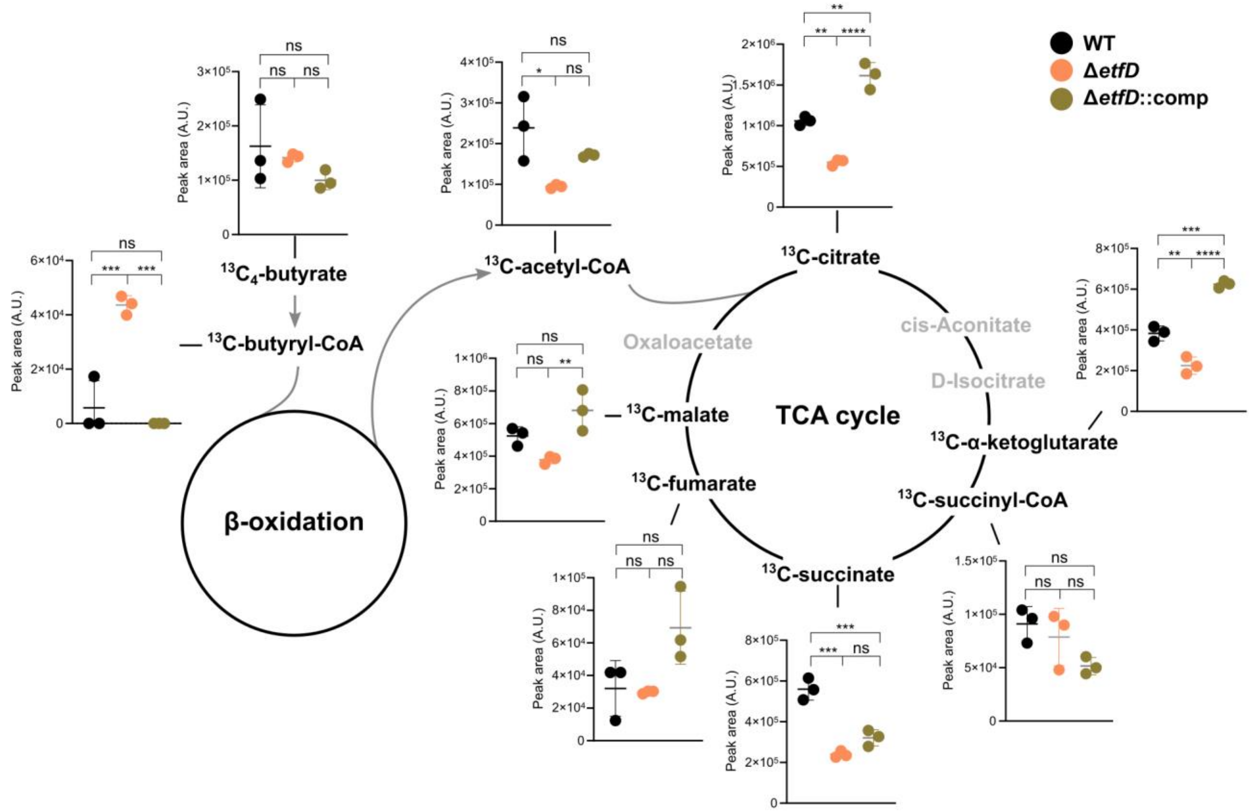


**Supplementary Fig. 5. Stable isotope tracing reveals a block in β-oxidation at the level of acyl-coA dehydrogenases.** Strains were grown on filters on top of solid medium permissible to Δ*etfD* growth for 7 days and then transferred to solid media with $^{13}C_4$-labelled butyric acid (2.5 mM) as single carbon source for 24 hrs. Levels of the indicated $^{13}C$-labeled metabolites (total $^{13}C$ pool, except for $^{13}C_4$-butyrate) were quantified by LC-MS analysis. Data correspond to 3 replicates of the second independent experiment. Error bars correspond to standard deviation. "Comp" stands for complemented. Statistical significance was assessed by one-way ANOVA followed by post hoc test (Tukey test; GraphPad Prism). *P < 0.05; **P < 0.01; ***P < 0.001; ****P < 0.0001. ns – not significant.

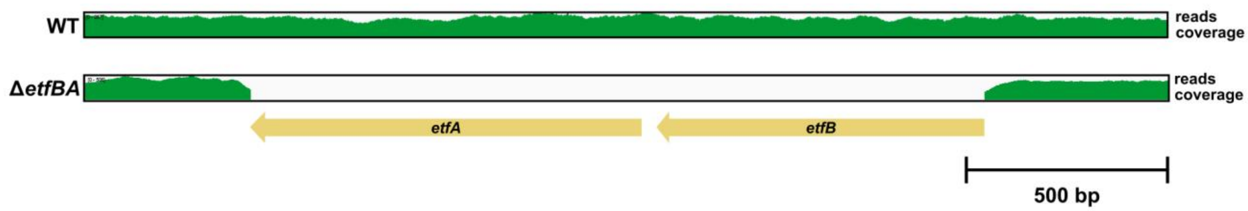


**Supplementary Fig. 6. Δ*etfBA* genetic identity confirmation.** Whole genome sequencing coverage showing the *etfBA* wild-type loci and gene deletion in Δ*etfBA*.

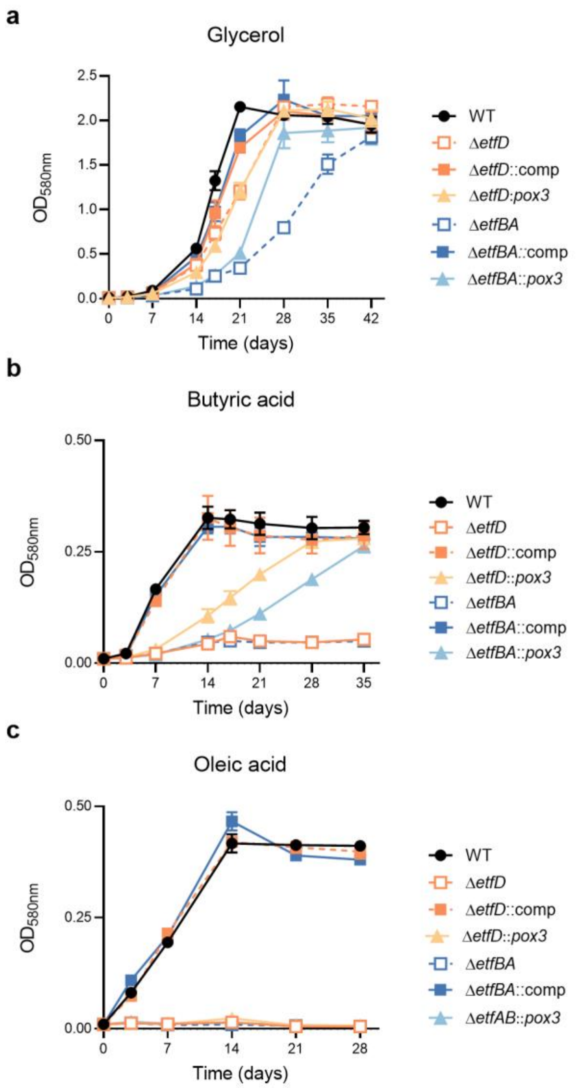


**Supplementary Fig. 7. Growth of Δ*etfD* and Δ*etfBA* expressing the acyl-CoA oxidase Pox3 in different carbon sources**. Strains were grown in media with 25 mM glycerol (**a**), 2.5 mM butyric acid **(b**), and 250 µM oleic acid (**c**). Oleic acid was replenished every 3 to 4 days to sustain growth and avoid toxicity. Data are averages of 3 replicates and are representative of 3 independent experiments. Error bars correspond to standard deviation. "Comp" stands for complemented.

**Supplementary Table 1.** ***Mycobacterium tuberculosis*** **whole genome sequencing analysis of Δ*etfD* and Δ*etfBA*.** This table presents the genetic polymorphisms present in the knockout strains in comparison with the parental strain.

| STRAIN | POSITION | GENE | DESCRIPTION | AA CHANGE | REF | ALT | # REF READS | # ALT READS | COVERAGE |
|---|---|---|---|---|---|---|---|---|---|
| **Δ*etfD*** | 132415 | *rv0109* | PE_PGRS1 | R346G | C | G | 1 | 411 | 412 |
| | 210094 | *rv0179c* | lprO | silent | A | G | 2 | 515 | 517 |
| | 528357 | NA | intergenic bet Rv0439c and Rv0440 | NA | C | A | 354 | 406 | 760 |
| | 837036 | *rv0746* | PE_PGRS9 | T445A | A | G | 0 | 406 | 406 |
| | 986207 | NA | intergenic bet Rv0887c and Rv0888 | NA | C | A | 0 | 531 | 531 |
| | 2874461 | *rv2553c* | Rv2553c | inframe ins V58 | G | GCCA | 5 | 256 | 261 |
| | 2933205 | *rv2604c* | snoP | D151E | G | T | 0 | 482 | 482 |
| | 3121725 | NA | intergenic bet Rv2813 and Rv2816c | NA | GGGTTTCCGTCCCCT CTCGGGGTTTTGGGT CTGACGACATGCTGA GCTGAGGCGCCGGAT GATGGTGGTGCTGAA | G | 15 | 193 | 208 |
| | 3961169 | *rv3523* | ltp3 | silent | G | C | 0 | 445 | 445 |
| **Δ*etfBA*** | 132415 | *rv0109* | PE_PGRS1 | R346G | C | G | 0 | 342 | 342 |
| | 210094 | *rv0179c* | lprO | silent | A | G | 3 | 368 | 371 |
| | 837036 | *rv0746* | PE_PGRS9 | T445A | A | G | 0 | 307 | 307 |
| | 986207 | NA | intergenic bet Rv0887c and Rv0888 | NA | C | A | 0 | 443 | 443 |
| | 2874461 | *rv2553c* | Rv2553c | inframe ins V58 | G | GCCA | 29 | 191 | 220 |
| | 2933205 | *rv2604c* | snoP | D151E | G | T | 0 | 326 | 326 |
| | 3121725 | NA | intergenic bet Rv2813 and Rv2816c | NA | GGGTTTCCGTCCCCT CTCGGGGTTTTGGGT CTGACGACATGCTGA GCTGAGGCGCCGGAT GATGGTGGTGCTGAA | G | 11 | 114 | 125 |
| | 4085080 | *rv3645* | Rv3645 | D307E | C | A | 5 | 398 | 403 |

Ref - Reference sequence

Alt - Alternative sequence

**Supplementary Table 2. EtfD interactors identified by mass spectrometry.**

| Rv | Gene | Description | EtfD-Flag : Control ratio |
|---|---|---|---|
| Rv0016c | pbpA | Probable penicillin-binding protein PbpA | 13.10 |
| Rv0050 | ponA1 | Probable bifunctional penicillin-binding protein 1A/1B PonA1 (murein polymerase) (PBP1): penicillin-insensitive transglycosylase (peptidoglycan TGASE) + penicillin-sensitive transpeptidase (DD-transpeptidase) | 14.74 |
| Rv0092 | ctpA | Cation transporter P-type ATPase a CtpA | 15.00 |
| Rv0202c | mmpL11 | Probable conserved transmembrane transport protein MmpL11 | 14.74 |
| Rv0338c | - | Probable iron-sulfur-binding reductase | 24.33 |
| Rv0496 | - | hypothetical protein | 11.50 |
| Rv0758 | phoR | Possible two component system response sensor kinase membrane associated PhoR | 18.00 |
| Rv0845 | - | Possible two component sensor kinase | 11.67 |
| Rv0862c | - | hypothetical protein | 12.22 |
| Rv0982 | mprB | Two component sensor kinase MprB | 16.67 |
| Rv1028c | kdpD | Probable sensor protein KdpD | 13.33 |
| Rv1179c | - | hypothetical protein | 12.57 |
| Rv1183 | mmpL10 | Probable conserved transmembrane transport protein MmpL10 | 10.00 |
| Rv1215c | - | hypothetical protein | 12.11 |
| Rv1265 | - | hypothetical protein | 11.60 |
| Rv1279 | - | Probable dehydrogenase FAD flavoprotein GMC oxidoreductase | 15.00 |
| Rv1281c | oppD | Probable oligopeptide-transport ATP-binding protein ABC transporter OppD | 12.11 |
| Rv1640c | lysX | Lysyl-tRNA synthetase 2 LysX | 16.25 |
| Rv1743 | pknE | Probable transmembrane serine/threonine-protein kinase E PknE (protein kinase E) (STPK E) | 10.33 |
| Rv1746 | pknF | Anchored-membrane serine/threonine-protein kinase PknF (protein kinase F) (STPK F) | 10.00 |
| Rv1747 | - | Probable conserved transmembrane ATP-binding protein ABC transporter | 24.83 |
| Rv1783 | eccC5 | ESX conserved component EccC5 ESX-5 type VII secretion system protein | 11.40 |
| Rv1797 | eccE5 | ESX conserved component EccE5 ESX-5 type VII secretion system protein Probable membrane protein | 14.33 |
| Rv1842c | - | hypothetical protein | 13.89 |
| Rv2051c | ppm1 | Polyprenol-monophosphomannose synthase Ppm1 | 11.00 |

| | | | |
|---|---|---|---|
| Rv2088 | pknJ | Transmembrane serine/threonine-protein kinase J PknJ (protein kinase J) (STPK J) | 11.58 |
| Rv2460c | clpP2 | Probable ATP-dependent CLP protease proteolytic subunit 2 ClpP2 (endopeptidase CLP 2) | 10.53 |
| Rv2703 | sigA | RNA polymerase sigma factor SigA (sigma-A) | 10.00 |
| Rv2933 | ppsC | Phenolpthiocerol synthesis type-I polyketide synthase PpsC | 13.89 |
| Rv2942 | mmpL7 | Conserved transmembrane transport protein MmpL7 | 10.00 |
| Rv2962c | - | Possible glycosyl transferase | 10.56 |
| Rv3029c | fixA | Probable electron transfer flavoprotein (beta-subunit) FixA (beta-ETF) (electron transfer flavoprotein small subunit) (ETFSS) | 11.60 |
| Rv3245c | mtrB | Two component sensory transduction histidine kinase MtrB | 17.37 |
| Rv3365c | - | hypothetical protein | 32.00 |
| Rv3519 | - | hypothetical protein | 12.22 |
| Rv3554 | fdxB | Possible electron transfer protein FdxB | 12.78 |
| Rv3610c | ftsH | Membrane-bound protease FtsH (cell division protein) | 12.40 |
| Rv3627c | - | hypothetical protein | 10.00 |
| Rv3693 | - | Possible conserved membrane protein | 10.56 |
| Rv3764c | tcrY | Possible two component sensor kinase TcrY | 16.11 |
| Rv3775 | lipE | Probable lipase LipE | 21.58 |
| Rv3779 | - | Probable conserved transmembrane protein alanine and leucine rich | 10.00 |
| Rv3794 | embA | Integral membrane indolylacetylinositol arabinosyltransferase EmbA (arabinosylindolylacetylinositol synthase) | 10.00 |
| Rv3808c | glfT2 | Bifunctional UDP-galactofuranosyl transferase GlfT2 | 11.67 |
| Rv3869 | eccB1 | ESX conserved component EccB1 ESX-1 type VII secretion system protein Possible membrane protein | 10.00 |
| Rv3876 | espI | ESX-1 secretion-associated protein EspI Conserved proline and alanine rich protein | 15.56 |
| Rv3885c | eccE2 | ESX conserved component EccE2 ESX-2 type VII secretion system protein Possible membrane protein | 11.03 |
| Rv3894c | eccC2 | ESX conserved component EccC2 ESX-2 type VII secretion system protein Possible membrane protein | 42.11 |
| Rv3895c | eccB2 | ESX conserved component EccB2 ESX-2 type VII secretion system protein Probable membrane protein | 11.67 |

**Supplementary Table 3. Strains used in this work.**

| Strain | Reference |
|---|---|
| *Mtb* H37Rv | Gift from C. Sassetti, University of Massachusetts. |
| *Mtb etfD*-tetOFF | This work |
| *Mtb* Δ*etfD* | This work |
| *Mtb* Δ*etfD* complemented | This work |
| *Mtb* Δ*etfD*::*pox3* | This work |
| *Mtb* Δ*etfBA* | This work |
| *Mtb* Δ*etfBA* complemented | This work |
| *Mtb* Δ*etfBA*::*pox3* | This work |
| *Mtb* H37Rv::Flag | [1] |
| *Mtb* Δ*etfD*::*etfD*-Flag | This work |

**Supplementary Table 4. Plasmids used in this work.**

| Plasmid | Reference |
|---|---|
| pNit-Rec-ET | [2] |
| pGMCgS-TetOff-13 | [3] |
| pMCK-phsp60-etfD | This work |
| pMCK-phsp60-etfBA | This work |
| pMCK-pTB38-pox3 | This work |
| pMEK-phsp60-etfD-flag | This work |
| pGMCK-hsp60-Flag | [1] |
